# The C-type lectin Schlaff ensures epidermal barrier compactness in *Drosophila*

**DOI:** 10.1101/381491

**Authors:** Renata Zuber, Khaleelulla Saheb Shaik, Frauke Meyer, Hsin-Nin Ho, Anna Speidel, Nicole Gerhing, Slawomir Bartoszewski, Heinz Schwarz, Bernard Moussian

**Affiliations:** Applied Zoology, Technical University of Dresden, Zellescher Weg 20b, 01217 Dresden, Germany; University of Tübingen, Interfaculty Institute of Cell Biology, Section Animal Genetics, Auf der Morgenstelle 15, 72076 Tübingen, Germany; Rzeszow University, Department of Biochemistry and Cell Biology, ul. Zelwerowicza 4, 35-601 Rzeszów, Poland; Max-Planck-Institut für Entwicklungsbiologie, Microscopy Unit, Spemannstr. 35, 72076 Tübingen; Université Côte d’Azur, CNRS, Inserm, Institute of Biology Valrose, Parc Valrose, 06108 Nice CEDEX 2, France

## Abstract

The stability of extracellular matrices is in general ensured by cross-linking of its components. Previously, we had shown that the integrity of the layered *Drosophila* cuticle relies on the presence of a covalent cuticular dityrosine network. Production and composition of this structure remained unstudied. In this work, we present our analyses of the *schlaff* (*slf*) gene coding for a C-type lectin that is needed for the adhesion between the horizontal cuticle layers. The Slf protein mainly localizes between the two layers called epicuticle and procuticle that separate from each other when the function of Slf is reduced or eliminated paralleling the phenotype of a cuticle with reduced extracellular dityrosine. Localisation of the dityrosinylated protein Resilin to the epicuticle-procuticle interface suggests that the dityrosine network mediates the adhesion of the epicuticle to the procuticle. Ultimately, compromised Slf function is associated with massive water loss. In summary, we propose that Slf is implied in the stabilisation of a dityrosine layer especially between the epicuticle and the procuticle that in turn constitutes an outward barrier against uncontrolled water flow.

**Summary statement:** Extracellular matrices adopt a stereotypic organisation for function during development. The lectin Schlaff assists adhesion reactions to ensure compactness of the epidermal cuticle in Drosophila.

## Introduction

Extracellular matrices (ECM) contribute to tissue shape and function. Their integrity depends on covalent and non-covalent interaction of their components. Collagen crosslinking in the articular cartilage by lysyl oxidases, for example, enhances tissue stability against physical wears (Saito and Marumo, 2010). Another prominent example is the apical extracellular layered network of lipids and proteins that constitutes the epidermal stratum corneum (Harding, 2004; Nishifuji and Yoon, 2013; Rogers et al., 1996). A defective stratum corneum in patients suffering different types of ichthyoses provokes a dry and scaly skin (Akiyama, 2017). Lamellar ichthyosis is caused by mutations in the gene encoding the central cross-linking enzyme transglutaminase that introduces covalent glutamine-lysine bonds. Extracellular dityrosine links catalysed by peroxidases have been identified in connective tissues and in response to oxidative stress (Keeley et al., 1969; Keeley and Labella, 1972; LaBella et al., 1967; Malencik and Anderson, 2003; Tenovuo and Paunio, 1979a; Tenovuo and Paunio, 1979b). While the molecular mechanisms and biochemical reactions of ECM network formation are well understood, the subcellular localisation of these processes are largely unexplored.

We address this issue by studying the molecular and cellular processes of insect cuticle differentiation. The insect cuticle is an ECM that consists of the polysaccharide chitin, proteins, catecholamines and lipids that interact with each other to form a layered structure including the outermost envelope, the middle epicuticle and the inner procuticle (Moussian, 2010; Moussian, 2013). It is produced and organised at the apical plasma membrane and in the region adjacent to it named the assembly zone. In the fruit fly *Drosophila melanogaster*, several proteins including Knickkopf (Knk), Obstractor A (Obst-A), Chitinase 2 and the chitin deacetylses Vermiform (Verm) and Serpentine (Serp) act in concert to ensure the stereotypic organisation of the larval cuticle during embryogenesis (Chaudhari et al., 2011; Moussian et al., 2006b; Pesch et al., 2015; Pesch et al., 2017). Stabilisation of the cuticle depends partly on a network of molecular bonds between different types of yet largely unknown proteins mediated by catecholamines, glutamine-lysine bridges and dityrosines (Shaik et al., 2012; Shibata et al., 2010; Wright, 1987). Catecholamine incorporation depends on a set of insect-specific enzymes including phenol-oxidases (sclerotisation) and occurs predominantly within the upper region of the procuticle called exocuticle. Glutamine-lysine crosslinking in *D. melanogaster* involves a transglutaminase that among others uses the chitin-binding proteins Cpr76Bd, Cpr47Ef, Cpr64Ac and Cpr97Eb as substrates suggesting that it acts within the procuticle. Dityrosine crosslinking was postulated to occur in the basal site of the procuticle adjacent to the apical plasma membrane of the epidermal cells (Shaik et al., 2012). Spatial information is, hence, available for these events. By contrast, the molecular and cellular mechanisms that control or mediate their localisation within the differentiating cuticle are unknown.

In this work, we have analysed the role of the C-type lectin Schlaff (Slf) in cuticle organisation and compactness in *D. melanogaster*. We demonstrate that Slf participates in the establishment of the dityrosine network within a distinct zone of the cuticle required for overall stability of the ECM.

## Results

### Cuticle phenotype of slf mutant larvae

Differentiation of the *D. melanogaster* larval cuticle is initiated at stage 15 of embryogenesis and ends shortly before hatching (Moussian et al., 2006a). The developing embryo and the ready-to-hatch larva almost fills the entire space of the egg (Fig. 1). Homozygous *slf* mutant embryos look normal when cuticle differentiation starts, but ready-to-hatch larvae retract from the egg-shell and the space between the larva and the egg-shell is filled with liquid (Fig. 1). When freed from the egg, the larvae contract and crumple and the cuticle occasionally detaches from the surface of the animal (Fig. S1). The head skeleton and the tracheae are, however, unaffected. When fixed with Hoyer’s medium, the cuticle detaches from the body surface and forms blisters (Fig.1). Larvae transheterozygous for an EMS-induced *slf* mutation and any deficiency uncovering the *slf* locus e.g. Df(2L)ED250 or Df(2L)BSC225 (see below) display the same phenotype as *slf* homozygous mutant larvae. Overall, the *slf* mutant phenotype is reminiscent of the phenotype caused by a deletion of the *alas* gene that codes for an enzyme of the heme biosynthesis pathway (Shaik et al., 2012).

**Fig. 1.**
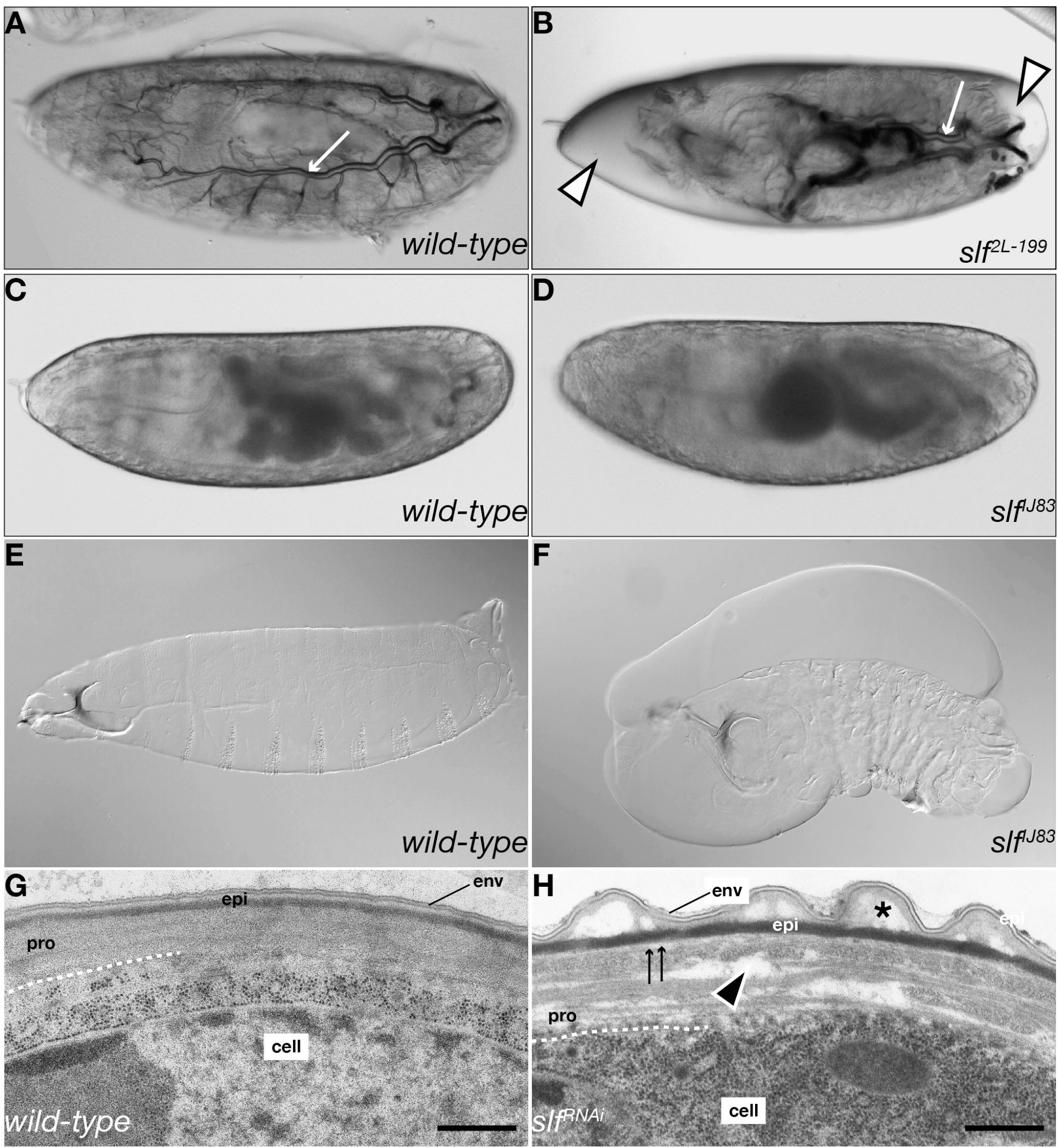
Homozygous slf mutant larvae contract and lose water whilst their cuticle detaches from the body surface. The ready-to-hatch living *wild type* larva fills the entire egg space (A), whilst the homozygous *slf* mutant larva (B) is separated from the egg case by liquid (white triangles). The head skeleton and the tracheae (white arrow) are unaffected in these larvae. The cuticle of the wild type larvae fixed in Hoyer’s medium acquires a spindle-like shape (C) whilst the cuticle of *slf* mutant larvae forms irregular bulges, especially in the head region, whereas the abdominal cuticle is crumpled (D). The cuticle of ready-to-hatch *wild type* larvae in transmission electron micrographs (E) is built of three distinct and tightly adhering layers: the external envelope (env) consisting of alternating electron-lucid and electron-dense sheets, the bipartite epicuticle and the chitinous procuticle (pro) that consists of chitin sheets (laminae). In *slf* mutant larvae (F) the procuticular laminal organization is disrupted by various sizes of electron-lucid regions (black triangle). Adhesion between the epicuticle and the procuticle is disrupted (arrows). The structure of the envelope seems to be normal but the envelope may detach from the epicuticle (asterisk).

For a detailed analysis of the cuticle phenotype, we examined the localisation of fluorescent-tagged cuticular proteins in wild-type and *slf* mutant larvae (Fig. 2). TweedleD-dsRed (TwdlD-dsRed) (Guan et al., 2006) and Cuticular Protein 67B-RFP (CPR67B-RFP) line the body surface of the wild-type larva. The region marked by TwdlD-dsRed detaches from the epidermis. By contrast, the region of CPR67B-RFP localisation does not detach and marks the body surface (Fig. 2).

**Fig. 2.**
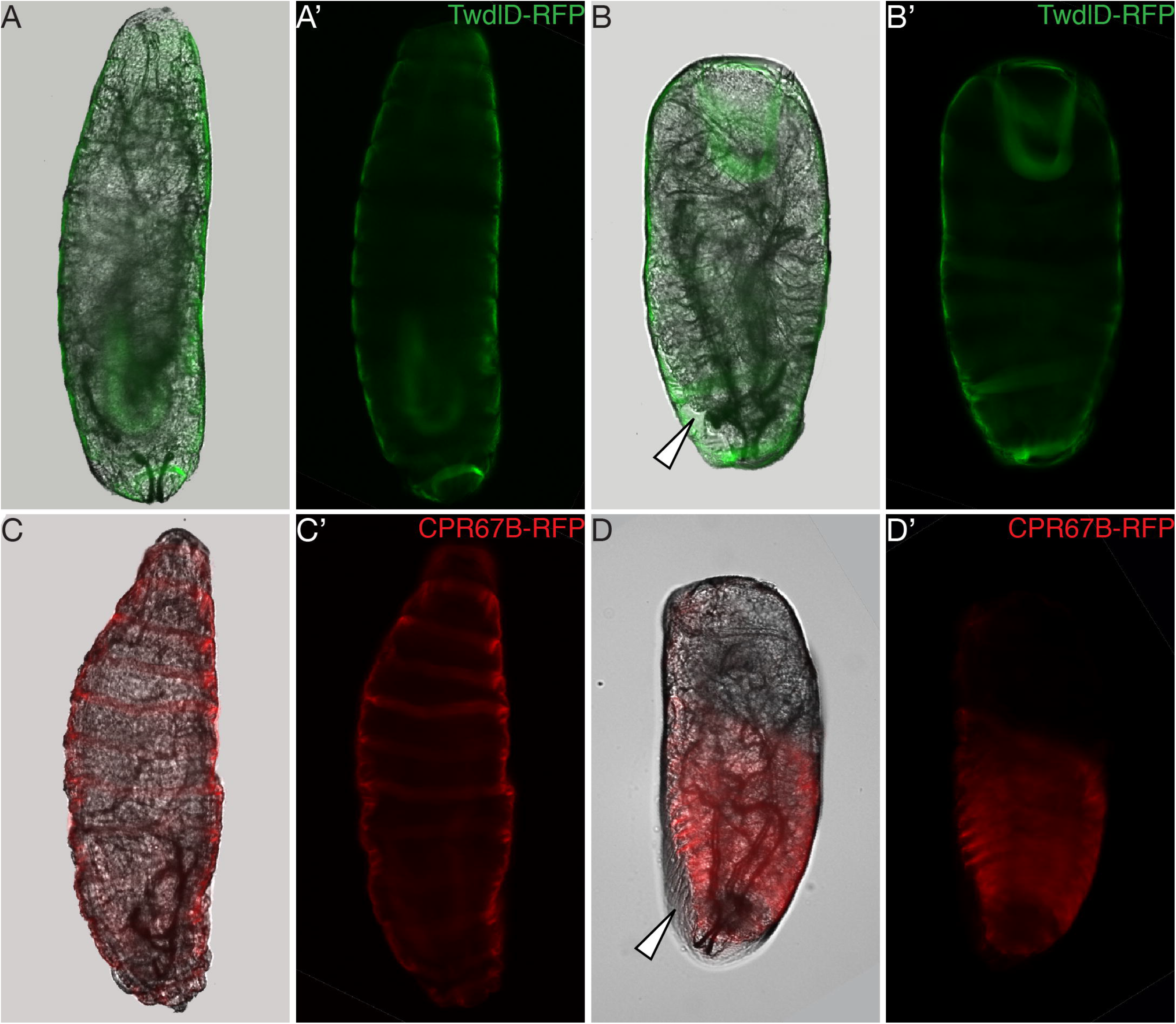
Cuticle of slf mutant larvae is delaminated. The layer of of TweedleD (TwdlD-dsRed, green) lines the body in living *wild type* larvae before hatching mounted in hydrocarbon medium (A, A’), whilst in *slf* homozygous mutant larvae this layer detaches from the body surface (B, B’, white arrow). The layer represented by CPR67b-RFP lines the body in *wild type* (C, C’) and *slf* (D, D’) mutant larvae, in spite of the detachment of the other cuticular layers in *slf* mutant larvae (white arrow).

Taken together, mutations in *slf* affect the integrity of the larval body. Especially, the adhesion between the TwdlD-dsRed and CPR67B-RFP domains depends on Slf. We assume that the observed liquid in the egg space of *slf* mutant embryos is the haemolymph that leaks out because through the loss of its integrity the cuticle has become permeable.

### Slf is not needed for the function of the inward barrier

To further inspect cuticle barrier integrity, we performed a dye penetration assay that we had developed recently (Wang et al., 2017a; Wang et al., 2016). Incubation of wild-type, *slf* and *alas* mutant ready-to-hatch embryos with bromophenol blue does not result in dye uptake, while *snurstorr snarlik* (*snsl*) mutant animals with a defective envelope (Zuber et al., 2018) do so (Fig. S2). Thus, Slf is not needed for protection of xenobiotic penetration through the cuticle.

### The cuticle of slf mutant embryos is delaminated

In order to understand the defects at the cellular level, we analysed the ultrastructure of the body cuticle of *slf* mutant larvae by transmission electron microscopy (Fig. 1). The wild-type body cuticle is composed of three biochemical distinct horizontal layers, the envelope, the epicuticle and the procuticle. The upper envelope consists of alternating electron-dense and electron-lucid films. The middle epicuticle is a bipartite matrix of cross-linked proteins and lipids. The procuticle contacting the apical surface of the epidermal cell is characterised by a helicoidal stack of chitin-protein sheets (laminae). In the cuticle of *slf* mutant ready-to-hatch larvae unstructured regions of various sizes disrupt the organisation of the laminae. The procuticle is occasionally separated from the above epicuticle. The upper tier of the epicuticle is not smooth. The envelope is continuous and its ultrastructure appears to be normal. In summary, cuticle compactness in *slf* mutant larvae is lost.

### Mutations in slf do not affect septate junctions

Loss of barrier function is observed in *Drosophila* embryos that have mutations in genes coding for septate junction (SJ) components (Izumi and Furuse, 2014). To test whether *slf* mutations affect SJ integrity, we investigated ultrastructure of the SJ in *slf* mutant embryos by transmission electron microscopy (Fig. S3). In the wild-type larva, SJs connect neighbouring epidermal cells. In the *slf* mutant larva the SJ ultrastructure is unchanged. The correct assembly of SJs does not exclude that they may have nevertheless lost their barrier function. We performed dye injection assays to analyse SJ barrier function (Fig. S3). When wild-type stage 16 embryos were injected with 10 and 3 kDa dye-conjugated dextran, the epidermal cells retained dextran within the body cavity. In *slf* mutant stage 16 embryos dextran retention is normal. Hence, we conclude that the loss of barrier function in *slf* mutant larvae is not due to defective SJ.

### Slf is a C-type lectin expressed in the epidermis

In order to understand the molecular defects caused by mutations in *slf*, we identified the gene affected by these mutations. The *slf* gene was initially mapped to the cytological position 25A to 25C on the left arm of chromosome 2 (see flybase.org). By deficiency mapping, we localised the mutations in an interval uncovered by Df(2L)BSC225 containing 10 loci. To narrow down the *slf* region, we attempted to reduce the number of candidate genes in a transgenic rescue experiment. Due to the cuticle defects observed in *slf* mutant larvae, we suspected that the factor affected might be associated with the apical plasma membrane or extracellular. A good candidate is CG3244 (Clect27) that was reported to be needed for wing cuticle integrity (Shibata et al., 2010). We recombined an insertion of the Pacman CHS322-140E11 (20233bp) that includes *CG3244* and the neighbouring gene *CG3294*, coding for a putative zinc-finger RNA-binding protein to the chromosome harbouring the *slf^2L-199^* mutation (Fig. 3). Homozygous *slf^2L-199^* larvae carrying the CHS322-140E11 insertion do not display the *slf* mutant phenotype. In *in situ* experiments, we detect the *CG3244* transcript in the developing epidermis during late embryogenesis when the cuticle is formed (Fig. 3).

**Fig. 3.**
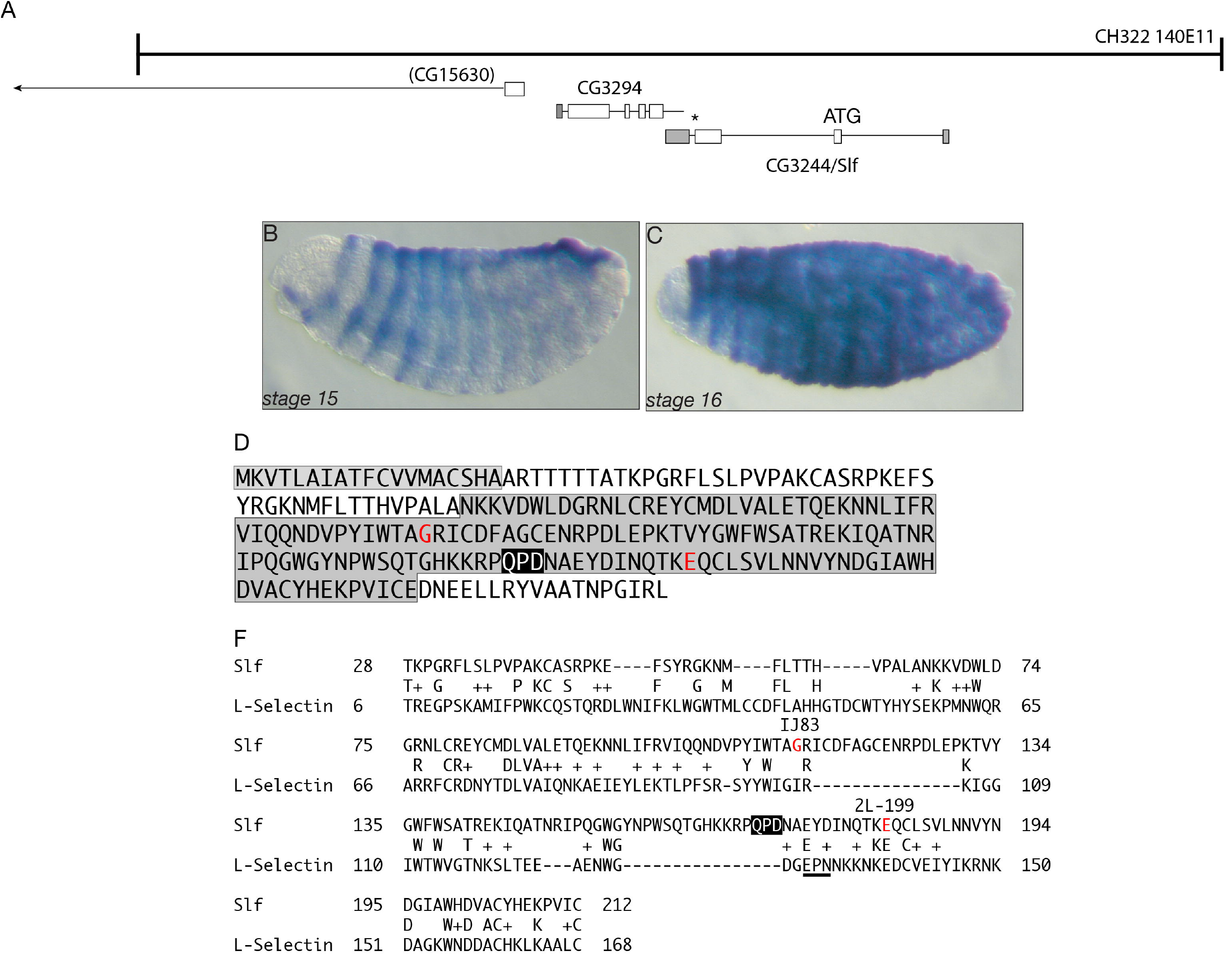
Slf is a C-type lectin. The CHS322-140E11 insertion including two genes (GC3294 and CG3244) recombined on the chromosome carrying the *slf 2L-199* mutation rescues the *slf* mutant phenotype (A). Detection of the CG3244 transcript in the developing embryos reveals the signal in the epidermis at stage 15 and 16 (B, C, blue). The Slf protein is composed of the N-terminal signal peptide (D, light grey) and the C-type lectin domain (D, dark grey) with the QPD motif (black), found in galactose binding lectins. In the sequence of *slf IJ83* mutants exchange of glycine to … has been found, whilst in the sequence of *slf 2L-199* mutants exchange of glutamate to … (D, F, red). There is weak homology between Slf and human L-Selectin (F).

According to the SignalP software, CG3244 possesses a signal peptide suggesting that it may be secreted. CG3244, a Ca^2+^-dependent lectin (C-type lectin), has recently been proposed to be a target of the transglutaminase that catalyses the cross-linking of proteins in the cuticle (Shibata et al., 2010). We sequenced the genomic DNA of the candidate CG3244 isolated from the two EMS alleles *slf^IJ83^* and *slf^2L-199^* and identified in each sequence a single point mutation that leads to an exchange of an amino acid (Fig. 3). These amino acids are highly conserved between CG3244 and homologous sequences. A rabbit antiserum produced against CG3244 failed to recognise an antigen in protein extracts from *slf* mutant larvae, while a 25 kDa protein was present in protein extracts from wild-type first instar larvae (data nit shown). Moreover, we were able to phenocopy the slf-mutant phenotype by RNA interference (RNAi) through the expression of UAS-driven *CG3244* RNA hairpin constructs in the epidermis (Suppl. Fig. S1). Thus, mutations in CG3244 are responsible for the *slf* mutant phenotype described above.

The Slf protein contains 231 amino acids and is composed of an N-terminal signal peptide and a C-type lectin domain (Fig. 3). The motif QPD especially within the C-type lectin domain is found in galactose binding lectins (Zelensky and Gready, 2005). Closely related sequences, probably Slf orthologs are found in other arthropods. A weak homology is detected to L-Selectins from vertebrates, which actually do not seem to have true counterparts in *Drosophila*. In order to determine the sugar moiety recognised and bound by Slf, we studied the binding capacity of Slf to mannose, galactose, lactose or N-acetyl-glucosamine (GlcNAc, the chitin monomer) in binding assays using agarose columns exposing the respective sugar. Slf extracted from stage 17 wild-type embryos was able to bind to mannose and galactose but not to lactose or GlcNAc (Fig. S4).

Taking all these data together, we conclude that *slf* encodes the C-type lectin CG3244, which potentially binds extracellular sugars but not chitin.

### Slf defines a new zone within the epidermal cuticle

Loss of cuticle compactness suggests that Slf is a coupling link between cuticle components. In order to examine the cuticular localisation of Slf we generated a C-terminally RFP-tagged Slf (Slf-RFP) version expressed in the larval epidermis under the control of the *tweedleM* promoter. To visualize the cuticle, we used a GFP-tagged version of the Tweedle-class protein Tubby (Tb-GFP) and an E-GFP-tagged version of the chitin-binding protein Obstructor (ObstE-GFP, (Tajiri et al., 2017)). Tb-GFP marks an apical region, while ObstE-GFP localises to a basal region adjacent to the epidermis. A Slf-RFP signal is detected in the whole procuticle with a strong signal in a thin region just below the Tb-GFP area and at the apical border of the ObstE-GFP layer (Fig. 4). Strong dots of an RFP signal occurred also under the procuticle, probably depicting intracellular vesicles. Thus, Slf localisation within the procuticle is necessary for cuticle compactness. We speculate that its accumulation in the apical region of the procuticle may define a new cuticle zone.

**Fig. 4.**
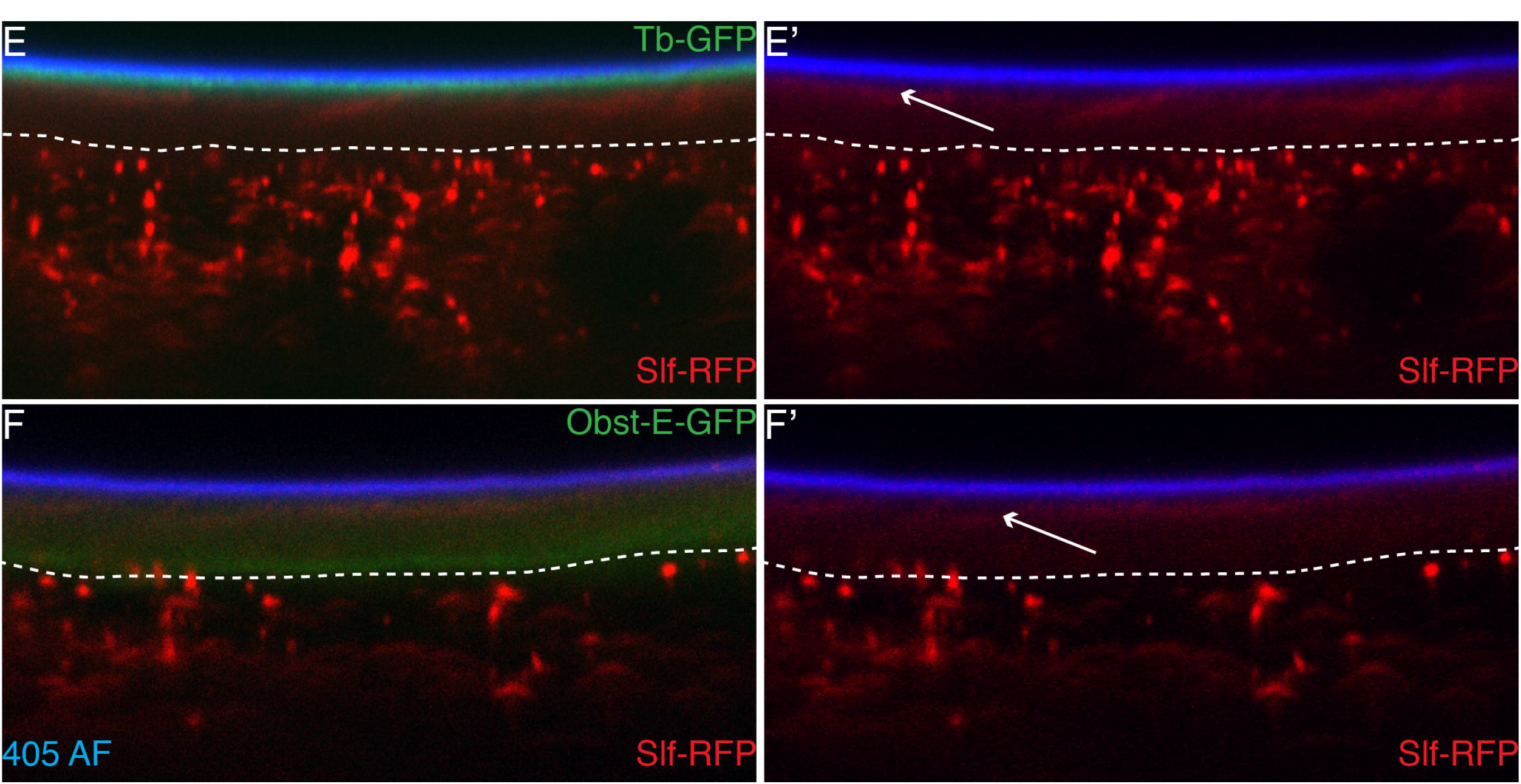
Slf protein localizes between the Tweedle layer and the procuticle. RFP-tagged Slf protein expressed in the epidermis under the control of the promoter of the *tweedleM* gene (*twdlm>slf-RFP*) forms particles in the cell and a thin layer in the cuticle of the living third instar larvae. The Slf layer (red) occurs below the Tweedle layer marked by a GFP-tagged Tubby protein, Tb-GFP (green; E, E’) and at the upper edge of the chitinous procuticle marked by GFP-tagged ObstE (green; F, F’). The 405-nm induced autofluorescence of the outermost cuticular layer envelope is marked in blue (E-F’).

### Slf is required for soft cuticle integrity

In *slf* mutant larvae, the soft body cuticle is disorganised, the head skeleton, by contrast, that consists of a melanised and hard cuticle is unaffected (Fig. 1). This observation suggests that Slf is needed especially in soft but not hard cuticle. To test this assumption, using the Flp/FRT technique (see Materials & Methods), we generated *slf* mutant clones in adult heads that are composed of hard sclerites connected by soft joints. Flies harbouring *slf* mutant tissue in the head fail to eclose and die within the pupal case. The overall anatomy of their head appears to be normal (Fig. 5). However, the ptilinum, a soft and elastic cuticle that expands to break open the pupa case, is ruptured. We thus reckon that soft cuticle integrity requires Slf function. To corroborate this interpretation, we down-regulated *slf* activity in the whole body of developing pupa by RNAi (Fig. S5). We observed that the cuticle in the leg joints, wing hinges, ventral abdomen and ptillinum were necrotic. The body parts with the hard cuticle appeared to be unaffected. These flies died in the pupal case or shortly after eclosion. In summary, our genetic experiments suggest that Slf is especially required in the unsclerotised, soft cuticle of larvae and adult animals.

**Fig. 5.**
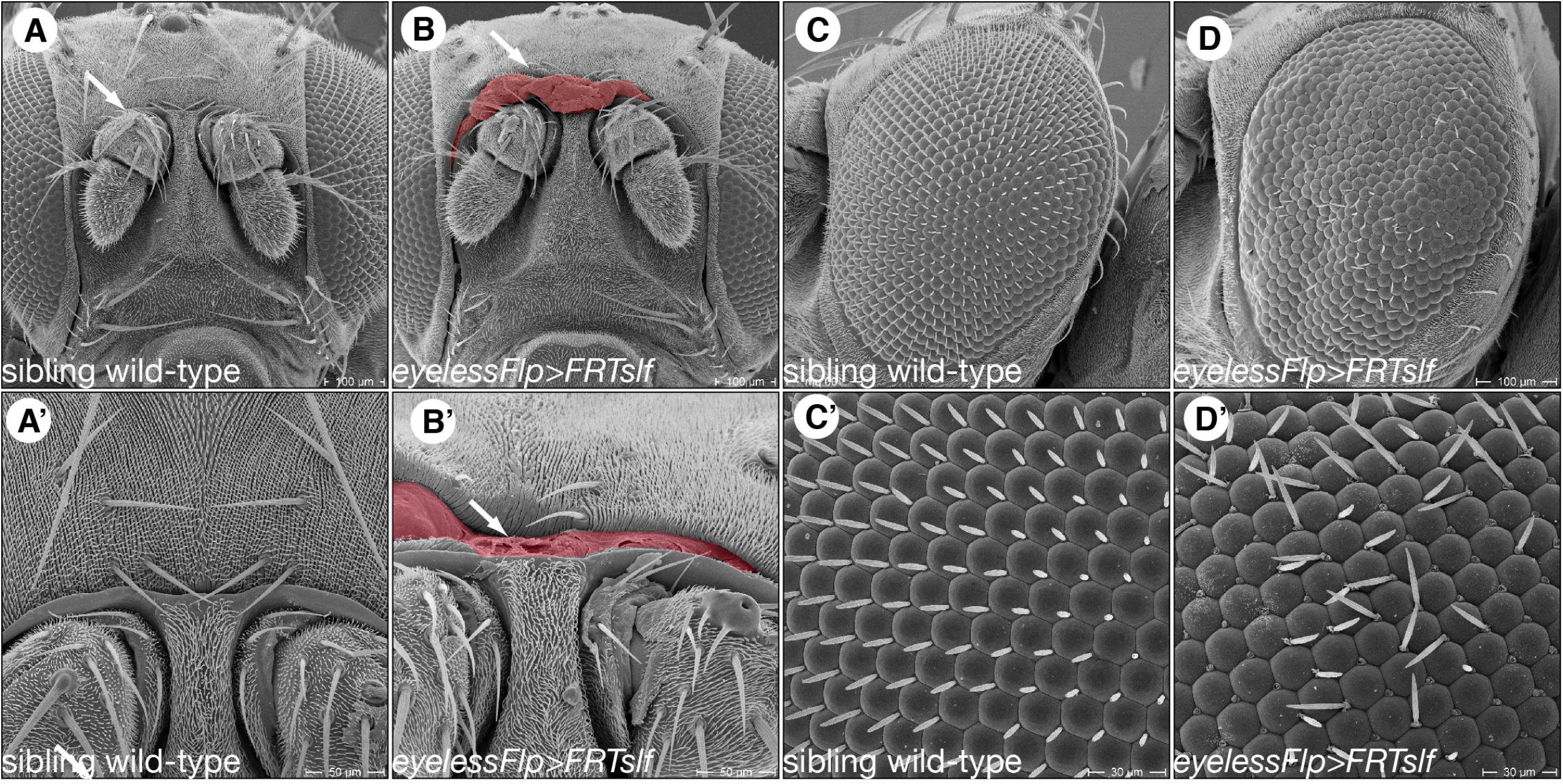
The ptilinum is disrupted in flies with down-regulated Slf. As shown in scanning electron micrographs, the head of the wild-type fly is composed of the large compound eye and sclerites bridged by rather narrow soft cuticular membranes that are not clearly exposed (A, A’). Homozygous *slf* mutant clones induced by the Flp/FRT system in the head of otherwise wild-type flies (*eylessFlp>FRT slf*) provoke disruption of especially the soft ptilinum at the forehead (arrow, B, B’) and the joining of the eye bristles with their basis (triangle, D, D’). The ommatidial structure is unchanged.

### Slf cooperates with heme synthesis pathway in dityrosine layer formation

Defects provoked by mutations in *slf* are reminiscent of those caused by mutation in *alas*, a gene encoding the delta-aminolevulinate synthase, which initiates the synthesis of heme (Figs. 1 & S1) (Shaik et al., 2012). Is there a genetic and molecular relationship between Slf and heme synthesis pathway? In order to answer this question, we performed a series of genetic and histological experiments. First, we examined embryos double-mutant for *alas* and *slf* mutations. The phenotype of these embryos was comparable to the ones provoked by mutations in either of the genes (Fig. S1). Assuming that both mutations represent loss-of-function situations, this observation suggests that these genes act in a common pathway. Consistently, reduction of larval *alas* or *slf* expression by RNAi caused a similar lethal phenotype (Fig. S6). Second, we tested whether Slf localisation may depend on Alas function. Using our anti-Slf specific antiserum, we find that Slf localises to the cuticle (Fig. 6). However, the thin L1 cuticle does not allow a more detailed localisation.

**Fig. 6.**
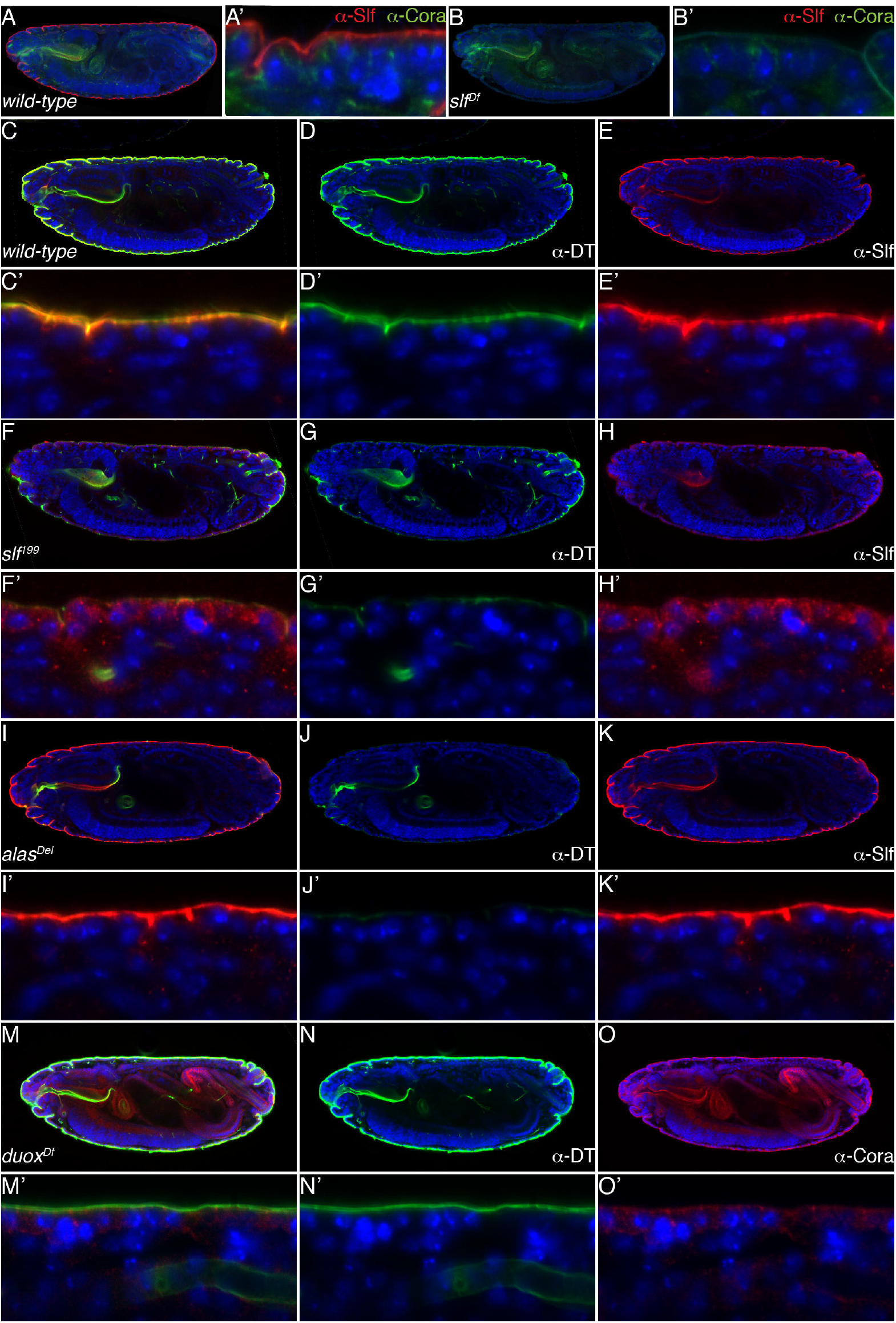
α-Slf antibody signal occurs in the cuticle of the embryos at late developmental stages. Probed with an α-Slf antibody (red), Slf is detected in the entire epidermal cuticle and the mouth hooks of *wild type* embryos at early stage 17 (A, A’), whilst embryos with a deleted *slf* gene do not show any α-Slf-signal (B, B’). Lateral boundaries between the cells are visualized by antibody staining against the junction protein Coracle (green) and the nuclei are visualized by blue DAPI staining (A-B’). The α-Slf antibody signal (red) co-localizes with the α-dityrosine antibody signal (α-DT, green) in the epidermal cuticle and the cuticle of the mouth hooks of the *wild-type* early stage 17 embryos (C-E’). The dityrosine signal occurs additionally in the tracheae where no Slf signal is present (C-E’, white arrow). In homozygous *slf^2L199^* mutant embryos at the early stage 17 the α-Slf signal (red) occurs inside the epidermal cells, whilst the α-DT signal (green) is strongly reduced in the epidermal cuticle, contrary to the mouth hooks and the tracheae, where it remains strong (F-H’). In early stage 17 embryos homozygous for the mutation in the *alas* gene the α-Slf signal (red) occurs in the epidermal cuticle and the mouth hooks, whilst the α-DT signal (green) is strongly reduced in the whole body (I-K’). In early stage 17 embryos with a deletion of the *dual oxidase* (*duox*) gene, the α-DT signal (green) is unchanged compared to *wild-type* embryos (M-O’). The lateral boundaries between the cells are visualized by α-Coracle staining (red, M-O’).

The phenotype of *alas* mutant larvae has been linked with the breakdown of the dityrosine barrier (Shaik et al., 2012). Using a DT specific antibody (α-DT), we tested whether the dityrosine network may depend on the presence of Slf. We observed that dityrosine signal intensity is reduced in these animals in the integumental cuticle, but not in the tracheal cuticle (not shown). This suggests that Slf might be either involved in dityrosine network formation or needed for the localisation i.e. stabilisation of dityrosinylated proteins to form a network. A well-known substrate protein modified by dityrosine links is Resilin (Andersen, 1964). We generated a Venus-tagged version of Resilin and co-expressed it with Slf-RFP in the cuticle of L3 larvae (Fig. 7). These chimeric proteins co-localise at the apical domain of Slf. Hence, Slf seems to be associated with dityrosinylated proteins. To further elucidate the relationship between Slf and Resilin, we expressed Resilin-Venus in third larvae with RNAi-induced reduced *slf* expression. We observed that Resilin-Venus is mislocalised in these larvae (Fig. 7). This suggests that Slf might be responsible for either the delivery or the stabilisation of dityrosine-forming proteins to the correct position in the cuticle.

**Fig. 7.**
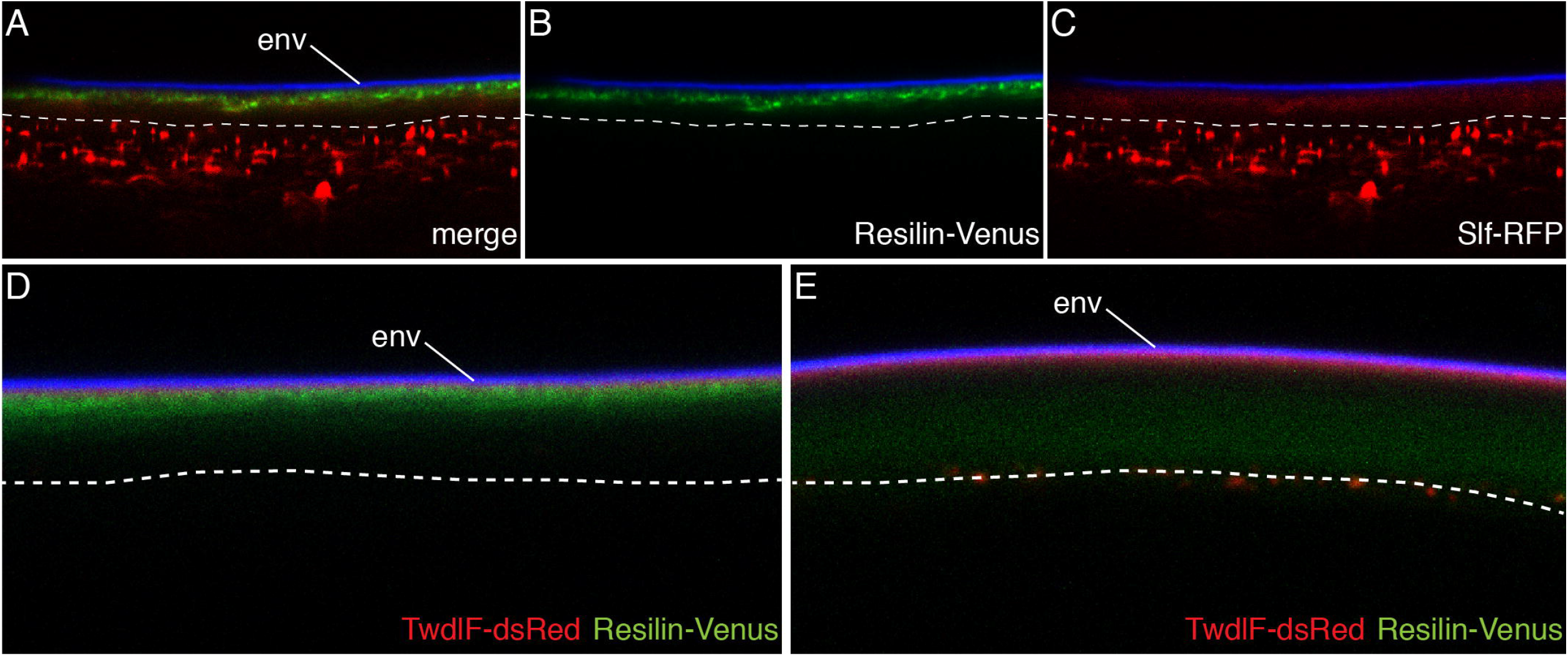
Resilin localization depends on Slf activity. In the cuticle of the third instar *Drosophila* larvae, the signal of RFP-conjugated Slf protein (A-A’’, red) overlaps with the signal of Venus-conjugated Resilin (A-A’’, green). The envelope blue signal shows auto-fluorescence of the envelope). In third instar larvae with down-regulated *slf* expression, the Resilin-Venus signal occurs in the whole procuticle (B’, green), whilst in the *wild type* larvae it is confined to a narrow region in the upper cuticle part (B). The localization of the TweedleF-dsRed protein is unchanged in *slf* RNAi third instar larvae (B’, red) in comparison to the *wild-type* larvae (B).

A well-known peroxidase involved in dityrosine formation in insects including fruit flies is the membrane-inserted Dual Oxidase Duox (Anh et al., 2011; Edens et al., 2001). Using a-DT specific antibody, we tested the presence of dityrosines in homozygous mutant embryos deficient for *duox*. In these animals, the dityrosine signal was comparable to the signal in wild type embryos (Fig.6). This suggests that either Duox is not involved in the formation of a larval dityrosine network, activity of maternally provided Duox is enough to catalyse the formation of the cuticle dityrosine network or another peroxidase may compensate decreased Duox activity.

Taken together, we conclude that Slf is a part of the dityrosine layer in the cuticle and the localisation of the dityrosinylated proteins to this layer depends on Slf activity.

### The Slf homologous CG6055 is needed for tracheal air-filling

Not only the epidermis, but also the tracheal tubes produce a cuticle that lines their surface (Moussian et al., 2006a). Slf is not present in the tracheal system (Fig. 3). According to the BDGP FlyExpress database (Konikoff et al., 2009), the two Slf homologous C-type lectins CG4115 and CG6055 are expressed in the tracheal tubes. To test whether these two factors may play a barrier role in the tracheae, we knocked-down the expression of CG4115 and CG6055 by RNAi. We expressed the respective UAS-driven stem loops by the tracheal-specific Gal4 driver btl-Gal4. RNAi provoked down-regulation of *CG6055* resulted in a subset of larvae that fail to air-fill their tracheae. Failure to air-fill has been repeatedly associated with loss of tracheal barrier function (Moussian et al., 2015; Tsarouhas et al., 2007; Wang et al., 2015). We therefore conclude that reduction of CG6055 function in the tracheae causes loss of barrier in the tracheal system.

In summary, Slf-like lectins are essential constituents of extracellular barriers in insects. Consistent with work on mammalian galectins (Argueso et al., 2009; Hikita et al., 2000), it is even thinkable that an important function of lectins in general is to contribute to the construction of epithelial barriers.

## Discussion

The insect cuticle is a water resistant barrier withstanding the internal hydrostatic pressure and preventing uncontrolled transpiration and water penetration. Previously, we had shown that a heme-dependent pathway is required to generate a dityrosine-based waterproof matrix within the cuticle of the *D. melanogaster* larva (Shaik et al., 2012). Recently, we reported on the role of the ABC transporter Snustorr (Snu) and the extracellular protein Snustorr-snarlik (Snsl) in the construction of an envelope-based anti-penetration and anit-transpiration barrier in *D. melanogaster* (Zuber et al., 2017). In the present work, we propose that the C-type lectin Slf cooperates with the heme-biosynthesis pathway to stabilise the distribution of the cuticle dityrosinylated proteins, exemplified by Resilin. The network of dityrosinylated proteins, in turn, is needed for correct contact between chitin laminae within the procuticle and between the procuticle and the epicuticle.

### Slf is a cuticular C-type lectin

Analyses of the Slf protein sequence suggest that it is a secreted galactose-binding C-type lectin. Our sugar binding data confirm the prediction that Slf is able to bind among others galactose. In *D. melanogaster*, galactose residues are found on side branches of N-glycans and on a tetrasaccharide that links glycosaminoglycans (GAGs) to serine residues of certain membrane-bound proteins such as glypicans and syndecans (Nakato and Li, 2016). Cuticle proteins have not been reported yet to harbour sugar moieties. Moreover, Slf is detected within the procuticle in *D. melanogaster* stage 17 embryos, especially accumulating at a distinct sheet at the apical border of the procuticle between the two zones marked by the cuticle proteins TwdlD and CPR67b. Based on these data, we assume that Slf is a cuticular C-type lectin contributing to late cuticle differentiation. Presumably, Slf exerts its function by binding an extracellular protein that carries a galactose. In principle, this finding is in line with data demonstrating the Slf (Clect27) is a cuticle protein that is essential for survival and needed for wing formation (Shibata et al., 2010). Moreover, it was shown that Slf is a substrate of the cross-linking enzyme transglutaminase that mediates covalent glutamine-lysine bonds. Down-regulation of transglutaminase expression, however, causes a mild cuticle phenotype compared to the strong *slf* mutant phenotype. Thus, taken together, Slf is a component of a composite extracellular network including non-essential covalent (glutamine-lysine bridges) and essential non-covalent (galactose binding) interactions.

We find that Slf is present in other insects. Thus, the role of Slf in the soft cuticle of other insects is probably conserved. According to information from the beetle base on the putative orthologue of Slf in the red flour beetle *Tribolium castaneum* (http://ibeetle-base.uni-goettingen.de/details/TC013911), injection of double-stranded RNA into larvae is 100% lethal. A phenotype has not been reported. However, this result underlines that Slf is also essential in other insects than *D. melanogaster*.

### Slf function is independent of the envelope

Classically, the outermost cuticle layer called envelope has been considered to be the bona fide desiccation barrier. In a recent work, we demonstrated that the extracellular protein Snsl and the ABC transporter Snu contribute to the establishment of the envelope in turn ensuring desiccation as well as penetration resistance (Zuber et al., 2017). The function of Snu is obviously conserved in other insects (Broehan et al., 2013; Yu et al., 2017). The envelope of *slf* mutant larvae is normal at the ultrastructural level. In addition, cuticle impermeability to xenobiotics is maintained in these larvae indicating that Slf is dispensable for an inward barrier. Furthermore, the procuticle is not disrupted in *snu* or *snsl* mutant larvae. Based on these evidences, we conclude that Slf and Snu/Snsl act in different pathways or mechanisms designed to establish a cuticular barrier preventing especially water loss.

### Slf is required especially in the soft unsclerotised cuticle

Elimination or reduction of Slf function especially affects the integrity of soft cuticle types including the larval body cuticle, the joint cuticle and the ptilinium. By contrast, hard cuticle types are largely unaffected. The major difference between hard and soft cuticles is the presence of an elaborate exocuticle in the hard cuticle that, as at the upper portion of the procuticle, consists of a sclerotised chitin-protein matrix. Based on this histological difference, we hypothesise that the region between the unsclerotised procuticle - called endocuticle in the hard cuticle - and the epicuticle is a region where components are cross-linked either by catecholamines (sclerotized exocuticle) or by dityrosines (soft cuticle). This region is apparently needed to prevent massive water loss through the cuticle.

### Slf is involved in organising cuticle compactness through production or stabilisation of the dityrosine network

Mutations in *slf* are embryonic lethal. Loss of Slf function entails massive water loss. By fluorescence microscopy, we show that the outer TwdlD-layer of the cuticle detaches from the inner CPR67b-layer of the cuticle in respective ready-to-hatch larvae. In addition, by transmission electron microscopy, we show that the procuticle of these larvae is loose. Thus, Slf is needed for compactness in the procuticle as well as the attachment of the TwdlD-to the CPR67b-layer within the cuticle.

The detachment of parts of the larval cuticle from the body is reminiscent of the *alas* mutant phenotype (Shaik et al., 2012). This suggests that Slf and Alas may contribute to the same structure in the cuticle. Alas is involved in the production of heme that is a co-factor of a yet unidentified oxidase catalysing the formation of a dityrosine network within the cuticle at the end of embryogenesis. We find that the cuticular dityrosine signal is reduced in *slf* mutant embryos and that the dityrosinylated cuticle protein Resilin is mislocalised in these animals. We conclude that Slf is required either for production or stabilisation of the dityrosine network that constitutes a barrier against water loss (Fig. 8).

**Fig. 8.**
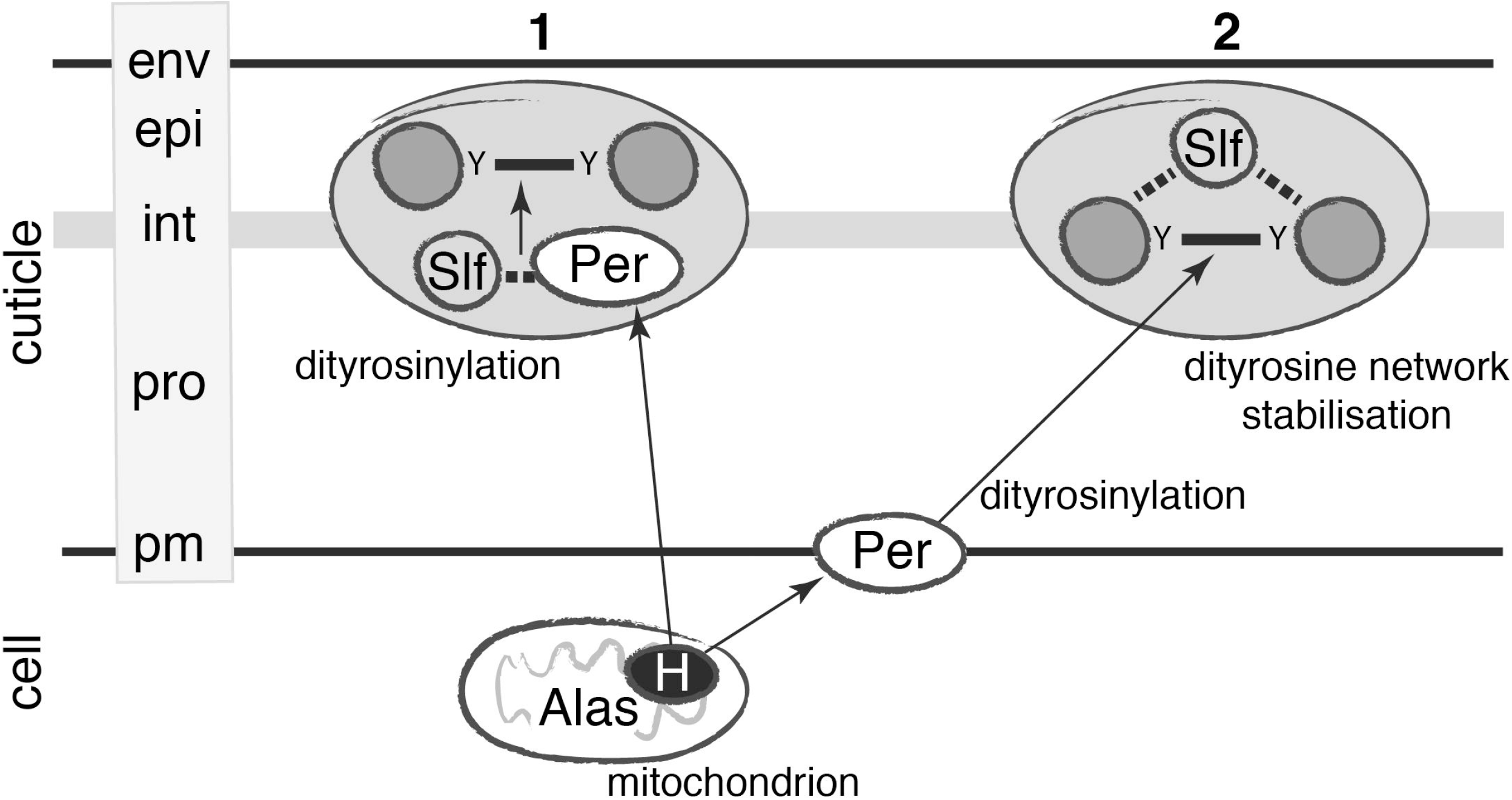
Slf promotes dityrosine formation or stabilises the diytrosine network. Our data allow proposing two alternative scenarios of Slf function. Either Slf assists directly a haem-dependent peroxidase (Per) at dityrosinylation of cuticle proteins such as Resilin (1), or it is needed to localize and stabilize the dityrosine network (solid lines) in the interface (int) between the epi- (epi) and procuticle (pro). Stabilization of the interface or interaction with a peroxidase may require sugar binding (dotted lines). The peroxidase may be inserted into the plasma membrane (pm) or extracellular; to simplify the scheme we have indicated only one possibility for each alternative scenario. Haem (H) is produced in the cytoplasm involving mitochondrial Alas. env envelope

Similarly, in vertebrates, galectin-3 forms an impermeable 500 nm thick lattice through the interaction with mucins at the surface of the ocular epithelium (Argueso et al., 2009). The presumed association of Slf with galactose-residues in a group of N-glycans or GAGs, its incorporation in a dityrosine and Gln-Lys network would in an analogous manner stabilise extracellular proteins required for cuticle integrity and barrier function. Slf is, hence, an adapter-like protein that glues different cuticular networks. Overall, we suspect that lectins may play a key role in ECM organisation.

## Materials & Methods

### Fly work and microscopy

Mutations were kept over balancers harbouring GFP or YFP constructs expressed under the control of either *Krüppel* or *Deformed*. This allows identification of homozygous mutant embryos, which lack any GFP expression. They were collected on apple juice agar plates garnished with a spot of yeast paste. For dextran injection experiments, embryos were dechorionated with bleach, and those of the desired stage were selected by hand, and dried on silica granulate for 4 minutes. 3kD dextran coupled to rhodamine and FITC-coupled 10kD dextran (Thermo-Fisher) were dissolved at a 10 mg/ml concentration in Sörensen injection buffer. For injections, these solutions were mixed at a 1:1 ratio, resulting in a 5 mg/ml concentration for each labelled dextran. Starting immediately after injection, behaviour of the fluorescence signal was monitored for about one hour using a Zeiss Axiophot microscope.

For cuticle preparations, larvae were deposited on a glass slide in Hoyer’s medium (Ashburner et al., 2005) covered by a coverslip and incubated at 65°C or 80°C overnight. They were examined by Nomarski microscopy on a Zeiss Axiophot microscope. Cuticle auto-fluorescence was examined after excitation with a 405nm laser on Zeiss LSM 710 or 880 confocal microscopes. For immunofluorescence microscopy, dechorionated embryos were fixed in Hepes buffered 3,7% formaldehyde for 20 minutes at room temperature, devitellinized and incubated with the respective antibodies, which were detected with appropriate secondary antibodies. Stained embryos were viewed on a Zeiss Axiophot, Olympus flowview FV1000, Zeiss LSM 710 or 880 confocal microscope. For permeability experiments, embryos were dechorionated, devitellinized and incubated in bromophenol blue solution following the protocol described in (Zuber et al., 2018). For electron microscopy, specimens were prepared following the protocol described in Moussian and Schwarz (Moussian and Schwarz, 2010). Samples for scanning electron microscopy (SEM) were prepared and analysed as published recently (Wang et al., 2017b). For live imaging, ready-to-hatch larvae carrying fluorescent cuticle markers were put on a glass slide into a drop of Halocarbon oil 700 (Sigma) and covered with a coverslip. Cuticle detachment was monitored using a Leica DMI8 fluorescent microscope. For live imaging of third instar larvae, larvae were anesthesized with ether, mounted in halocarbon oil on a glass slide and covered with a coverslip. Fluorescence was observed on a Zeiss LSM 880 microscope. Images were prepared using Adobe Photoshop and Illustrator CS6 software.

### Generation of homozygous slf mutant clones

The *slf^2L199^* allele was induced on a chromosome carrying FRT (Flipase Recognition Target) sequence (Luschnig et al., 2004). These flies were crossed to flies carrying a lethal mutation on a 2nd chromosome with FRT sequence and expressing Flipase in head driven by the *eyeless* promoter (*eye>flipase*). The progeny carried *slf*, FRT on one second chromosome, FRT on another homologous second chromosome and expressed Flipase in head of developing flies. As a consequence of the Flipase activity, *slf* homozygous clones were generated in the head of developing pupae.

### RNA interference

To generate flies expressing hairpin RNA against *slf* (*slf^RNAi^*) in the epidermis of pupae the UAS/Gal4 system was used (Brand and Perrimon, 1993). Flies carrying *slf^RNAi^* under the control of the UAS promoter (*UAS>RNAi-slf*, from NIG-Fly, Kyoto, Japan) were crossed with flies harbouring *Gal4* under the control of the promoter of the *knickkopf* gene (*knk>gal4*). The progeny eclosing of the pupae was observed. RNAi experiments to suppress the expression of *CG4115* and *CG6055* were conducted using appropriate UAS-driven hairpin constructs (Dietzl et al., 2007) under the control of the tracheal-specific driver *btl*-Gal4.

### Molecular Biology

Standard molecular techniques (PCR, sequencing) were applied to identify and characterise the *slf* gene as presented in figure 3.

### Carbohydrate binding assays

Proteins from 60 ready-to-hatch embryos were extracted in TCS buffer (10mM Tris-HCl, 10mM CaCl2, 150mM NaCl, ph 7.4). Protein extracts were incubated with D-Galcatose agarose (Thermo), Lactose-, Mannose-or N-acetyl-glucosamine sepharose (all three GALAB) beads. Western blotting and immune-detection was performed as previously described (Norum et al., 2010).

## Author contributions

RZ, KSS, FM, HH, AS, NG performed the experiments. RZ, SB, HS and BM analysed data. RZ and BM wrote the manuscript.

---

*Fig. S1 Dityrosine distribution depends on Slf but not on Duox*

The *wild-type* ready-to-hatch living larva fills the entire egg (A). Ready-to-hatch larvae with eliminated or reduced *slf* function (*slf^IJ83^, slf^2L-199^, slf deficiency, slf^RNAi^*) are contracted and the space between the embryo and the egg case is filled with liquid (B-E). Their tracheal system is air-filled and the head skeleton seems to be unaffected. The phenotype of the homozygous mutants carrying loss-of-function insertion in the *alas* (F) gene is reminiscent of the *slf* phenotype, but, additionally the tracheae are not air-filled. The phenotype of the *slf alas* double mutant larva resembles *alas* mutant embryos (G). The tracheae of the homozygous mutants in the *dual oxidase* (*duox*) gene are not air-filled, but the larvae do not contract in the egg (H).

After freeing from the egg and keeping the living larvae in halocarbon oil under the coverslip, the *wild type* larvae larvae stretch and their cuticle lines the body surface (I), whilst the cuticle of the *slf* mutant (J,K), *alas* mutant (L) and the *slf, alas* double mutant (M) larvae to a lesser or greater extent detaches from the body surface. In *duox* mutant larvae only a thin layer of the cuticle, probably the envelope detaches from the surface (N).

In Hoyer’s cuticle preparations, the envelope as visualized by a 405nm laser (blue) lines the body surface of the wild-type larva (O). In *slf* (P, Q) and *alas* (R) mutant larvae, the envelope forms small blisters at the ventral (vs) and large blisters at the dorsal side (ds) of the body. In *slf, alas* double mutant larvae, it detaches from the whole body forming large blisters (S). In *duox* mutants it forms small blisters on the ventral side of the body only (T).

*Fig. S2 Slf is not needed for inward barrier function*

The cuticle of the living *wild-type, slf, alas* and *duox* homozygous mutant ready-to-hatch larvae is impermeable for bromophenol blue (bpb). The upper panel shows the larvae before incubation and the lower panel after incubation with bpb. Larvae homozygous mutant for *snsl* exhibiting a defective envelope are permeable for bpb, which leaks into the larvae and stains them with a dark blue colour.

*Fig. S3 Septate juctions of the slf mutant larvae are normal*

Septate junctions (SJ) connect neighbouring epidermal cells as shown in electron micrographs of late *wild type* embryos (A). Comparably, septate junctions of the *slf* mutant larvae are unchanged (B). In larvae carrying mutation in gene encoding septate junction component Coracle, the septate junctions are not present (C). Epidermal cells of stage 16 *wild type* (D) or *slf* mutant embryos (E) contain particles of 10 kDa dye-conjugated dextran that was injected in their haemolymph. to the retains in the body cavity (asterisk; cut: cuticle, ec: epidermal cells), By contrast, in stage 16 coracle mutant embryos (F), the dextran signal is also detected in lines probably representing the lateral membrane (triangle).

*Fig. S4 Slf protein binds in vitro to galactose and mannose, but not to GlcNAc and lactose*.

Protein extracts from the wild type larvae before hatching show distinct band in size 27kDa representing the Slf protein (A, B). This band is missing in protein extracts from the *slf* deficient larvae (A, ?). A band in this size is visible in the eluates of the mannose (A) and galactose (B) columns, whilst not present in the eluates of the GlcNAc and lactose (A) columns.

*Fig. S5 Flies with down-regulated Slf activity show soft cuticle damages and necrosis*

By light microscopy, in flies with down-regulated Slf activity by RNAi, we observe necrosis in the soft cuticle regions like the dorsal abdominal region (asterisk, F), at joints (arrow, H) and the wing hinge (arrowhead, J) in comparison to the intact cuticle in *wild-type* flies (E, G and I).

*Fig. S6 Down-regulation of slf or alas causes a similar larval phenotype*

The wild-type L1 and L3 larvae have a slender body shape (A,D). L3 and L1 larvae with down-regulated *slf* expression in the epidermis and tracheae or down-regulated ubiquitous *alas* expression induced by RNAi (*slf^RNAi^*; *knk*-Gal4 and *alas^RNAi^*; *L370*-Gal4, respectively) are podgy and have melanised injuries on their surface (arrows, B,C).

We were unable to compare the same larval stages for down-regulation of *slf* or *alas*, because *slf^RNAi^*; *L370*-Gal4 larvae die within the egg case showing the *slf* mutant phenotype, and *alas^RNAi^*; *knk*-Gal4 do not show any phenotype.

